# A Waddington Epigenetic Landscape for the *C. elegans* embryo

**DOI:** 10.1101/2020.01.01.892414

**Authors:** Ahmed Elewa

## Abstract

Waddington’s Epigenetic Landscape provides a visual model for both robust and adaptable development. Generating and exploring a Waddington epigenetic landscape for the early *C. elegans* embryo suggests that the key shapers of the landscape are genes that lie at the nexus between stress response and behavior and include genes that are regulated by transgenerational neuronal small RNAs. Curiously, several genes shape the early landscape of one lineage and then pattern, differentiate or are enriched in another lineage. Additionally, paralogs with similar expression profiles contribute differently to shaping the modeled landscape. This work suggests that robust embryonic development is initialized by differential deployment of redundant genes and by transgenerational cues that configure the epigenetic landscape to adapt to a changing world.

## Introduction

Development balances consistency and plasticity. The consistency of development is demonstrated by its robustness against perturbations (Félix and Wagner, 2008; Wagner, 2005). A key cause for robustness is redundancy, multiple alternative paths that lead to the same destination, so that if one fails, other paths provide way (Maduro, 2015; Motegi and Seydoux, 2013; Ritter et al., 2013). At the same time, development cannot afford rigidity. In fact, the very redundancy that provides robustness also allows sampling alternative paths, for although they all may lead to the same place, different circumstances may favor different routes (Barghi et al., 2019; Natarajan et al., 2016). Development therefore relies on a multiplicity of paths that provide robustness on one level and flexibility on another. But how does a developing system decide which path to take? On a population level, a species may afford trial and error, but on an individual level, finding the optimal path forward from the first try is a challenge of existential proportions. Understanding how the decision is made on an individual level requires understanding how robustness is set up and how flexibility is concealed within.

To picture how robustness is built into development Conrad Waddington created a visual metaphor known today as Waddington’s Epigenetic Landscape (Waddington, 1957). The landscape captures two key features of development, first that the end products of development are sharply distinct (nerve, skin, muscle, etc.) and second that the process is robust against disturbances. To illustrate these two features, the landscape depicts a declining surface representing developmental time on one axis and the end products of development represented as valleys across the perpendicular axis (Figure 1A). The depressions (or canals) leading to the end products convey the notion of canalization, the resilient trajectory of development, so that if a ball representing a developing system were to roll down it would be confined to canals despite external disturbances. A basis on which canals are formed in the landscape was also conceived by Waddington, and involves pegs representing genes pulling on the surface from beneath by a network of ropes with tensions representing the chemical forces exerted by gene products (Figure 1B). Therefore, according to Waddington’s model, gene activity shapes the titled epigenetic landscape and creates canals; trajectories of fate that cells of an organism are more likely to traverse during development even when challenged with perturbations.

**Figure 1.**
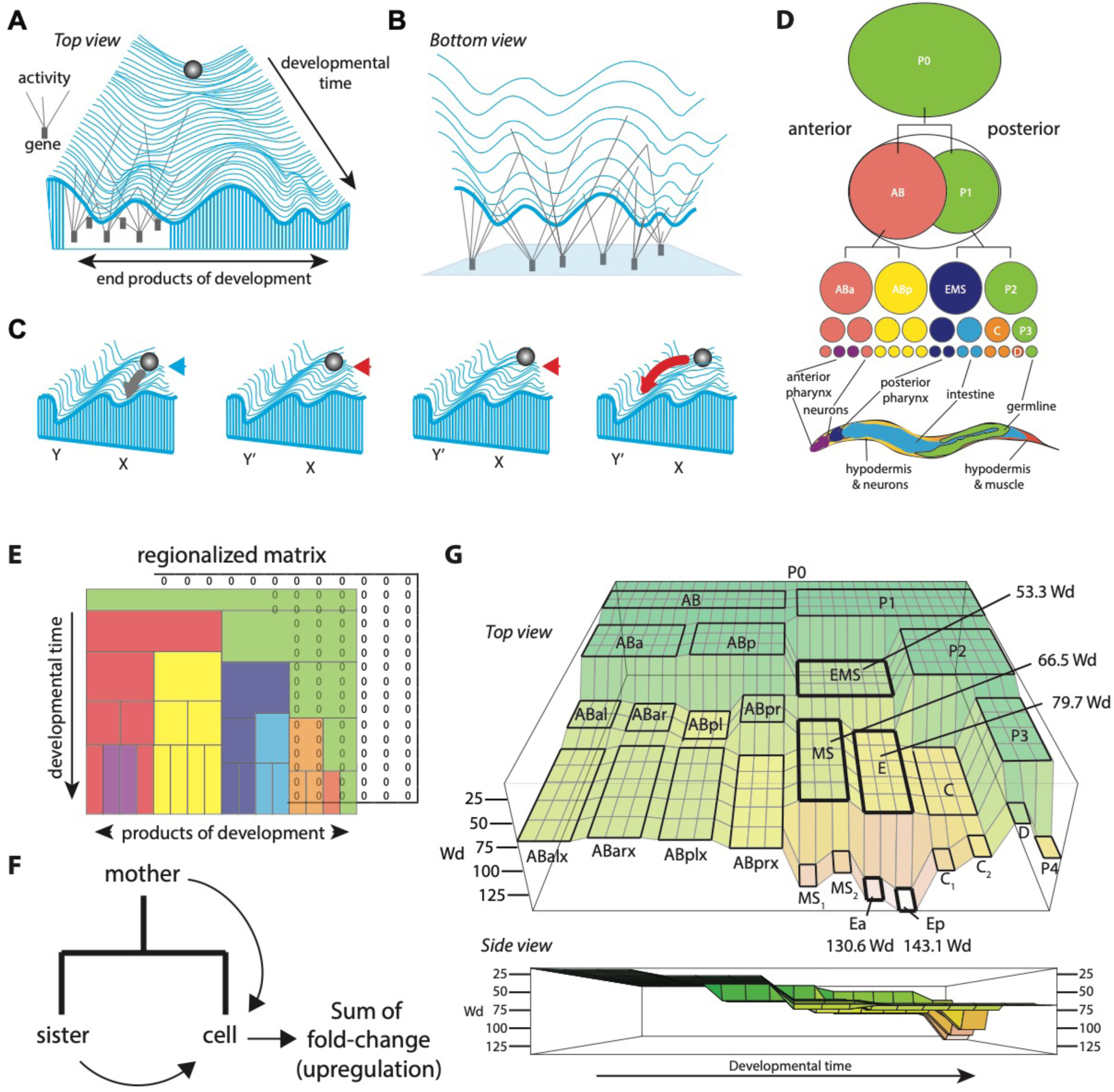
A Waddington Epigenetic Landscape for *C. elegans* early embryogenesis. **A)** A recreation of Waddington’s Epigenetic Landscape. **B)** A bottom view of the landscape depicting the effect of gene activity on canalization. **C)** A recreation of an example for organic selection from Waddington’s “Strategy of Genes”. A main valley leads to the character X and a side branch to Y. The developing tissue does not get into the Y path unless an environmental stimulus (red arrow) pushes it over the threshold. However, a mutant allele (red arrow) can lower the threshold to some extent, or cause the threshold to disappear. When this allele is selected for, the side branch Y’ will be the default, even when in the absence of an environmental stimulus. **D)** *C. elegans* early embryogenesis. Invariant cleavages lead to lineages that give rise distinct body parts. **E)** The lineage of early *C. elegans* embryogenesis converted to a regionalized matrix. Depicted in colors are regions corresponding to specific cells (compare to Figure 1D). Also depicted is an initial matrix of zeros. **F)** Epigenetic tension is calculated as the sum of upregulation in a cell compared to sister and to mother. **G)** The epigenetic landscape of *C. elegans* early embryogenesis computed from the Tintori *et al*. dataset. Depicted are regions for each cell and epigenetic tension values for EMS to E daughter cells. Colors represent elevation (terrain). Side view highlights the overall downward tilt of the landscape.

In addition to offering a visual metaphor for robustness that endures till today, Waddington’s legacy also includes a theory for developmental plasticity and adaptation. A question that Waddington addressed was how physiological adaptations become genetically fixed so that when the circumstances that triggered those adaptations no longer exist, the adaptations still appear (Waddington, 1957). Using the epigenetic landscape for visual explanation and by integrating a notion of organic selection developed by James Baldwin, Waddington envisaged how existing genetic variants that are more adaptive to new circumstances are selected for in a population (Crispo, 2007; Waddington, 1957). With natural selection, these genetic variants will mold the epigenetic landscape and canalize the adaptation so that even if the circumstances no longer exist, the adaptation would still appear (Figure 1C). According to this view, genetic redundancy offers a variety of alternatives, some of which may prove more adept to overcome new challenges. These advantageous varieties spread to subsequent generations and shape the epigenetic landscape in a population so that what was once an alternative becomes the robust course of development. However, with the discovery of RNAi and the inheritance of endogenous small RNAs and their accompanying protein argonautes, an alternative path to inherited adaptations has been uncovered (Buckley et al., 2012; Fire et al., 1998). These forms of transgenerational epigenetic inheritance provide the benefits of an adaptation to individuals before it is genetically fixed in a population (Buckley et al., 2012; Fire et al., 1998; Moore et al., 2019; Posner et al., 2019).

With only 302 neurons, *C. elegans* is able to navigate complex environments in search of food, safety and sex. Upon noticing that an attractive source of nutrition is in fact harmful (e.g. *Pseudomonas aeruginosa* – PA14), worms adapt by developing and remembering an aversion to food (Zhang et al., 2005). Moreover, this behavioral adaptation can be transmitted from mothers to their progeny in a manner dependent on the germline argonaute Piwi/PRG-1 and its piRNAs (Moore et al., 2019). PRG-1 and piRNAs are necessary for germline development and guard its integrity by silencing of transposable elements and foreign nucleic acids (Bagijn et al., 2012; Batista et al., 2008; Ghildiyal and Zamore, 2009; Lee et al., 2012; Shirayama et al., 2012). The silencing initiated by PRG-1 and piRNAs can be transferred to subsequent generations and maintained by <u>w</u>orm specific <u>a</u>r<u>go</u>nautes (WAGOs) such as WAGO-1 and WAGO-9 and a separate class of small RNAs known as 22Gs (Ashe et al., 2012; Gu et al., 2009; Yigit et al., 2006). WAGOs are responsible for the heritable chromatin modifications (e.g. H3K9me3) that ensure silencing for several generations (Buckley et al., 2012). WAGO-9, also known as HRDE-1, is responsible for the inheritance of a class of 22Gs that are generated in neurons and transmitted to the germline and which can later control behavior for multiple generations (Buckley et al., 2012; Juang et al., 2013; Posner et al., 2019). Therefore, endogenous small RNA pathways provide mechanisms for behavioral adaptations to be communicated to posterity without the need for the slower process of genetic assimilation.

Countering the silencing effect of PRG-1 and WAGOs, the argonaute CSR-1 acts in the germline to ensure the transcription of genes that should be expressed (Claycomb et al., 2009). Additionally, CSR-1 acts to tune maternal transcripts loaded in oocytes to ensure robust early *C. elegans* embryogenesis (Gerson-Gurwitz et al., 2016). However, it remains unclear how the aforementioned small RNA pathways maintain the robustness of germline and embryo development and at the same time act as agents for adaptive plasticity. Moreover, we do not know how the information in small RNAs is converted once again to behavior. Here I generate a Waddington epigenetic landscape for early *C. elegans* embryogenesis to explore how robustness and plasticity are expressed at the beginning of development. Surprisingly, the entry points of canals leading to robust and discrete developmental end products are initially shaped, not by master regulators of development, but by genes influenced by physiological adaptations and include genes identified as contributors to transgenerational behavior.

## Results

### Generating a Waddington epigenetic landscape from single-cell RNAseq data

A surface resembling a Waddington epigenetic landscape should satisfy three conditions. The surface should have a downward tilt, should display canals that lead to the end products of development and the curvature of the surface should be determined by gene activity. To generate a surface that meets these conditions, I took advantage of the properties inherent in *C. elegans* early embryogenesis and a published dataset that profiled the transcriptomes of early embryonic cells at single-cell resolution (Sulston et al., 1983; Tintori et al., 2016). *C. elegans* embryogenesis commences with a fixed series of cell cleavages that generate blastomeres, which then give rise to lineages that build the body (Figure 1D). Converting the lineage to a regionalized matrix results in a two-dimensional surface with developmental time on one axis and on the second axis, the blastomeres and their lineages that lead to the end products of development (Figure 1E). The matrix is regionalized so that each cell in the lineage has a set of points corresponding to its developmental time and proportionate size. Since early *C. elegans* embryogenesis occurs through cleavages, this means that daughter cells are smaller than their cleaved mothers (Figure 1E).

Next, by adopting gene expression as an approximation for gene activity, I computed a value that corresponds to “rope tension” from each gene to each region in the matrix. If the tension were based on gene expression measured in transcripts per million (TPM), then each gene would pull on each region in the matrix with a tension equal to its expression level. However, with gene expression from each cell normalized to 1 million transcripts, the overall effect will be the same in each region, leading to flat surface. Since the aim is to create a landscape that communicates canalization, another approach is necessary. Instead of expression level, rope tension is calculated from gene upregulation (**Methods**). Comparing gene expression between each cell and its mother and sister, and summing the fold changes in all upregulated genes leads to a quantity equal to the extent of upregulation in a cell vis-à-vis its mother and sister. I refer to this quantity, the sum of all upregulations in a cell, as “epigenetic tension”, which I measured in waddings (wd) (**Figure 1F and Methods**).

Visualizing the lineage matrix up to the 16-cell embryo after populating each region with its epigenetic tension value results in a surface with a downward tilt and rudiments of canals appearing at the beginning of each lineage. The downward tilt is due to the overall increase in gene upregulation during blastomere specification as a transcriptome departs from pluripotent uniformity to fate determination. For instance, epigenetic tension at the region corresponding to the blastomere EMS equals 53325.37 wd, or 53.3 Waddington (1000 wadding = 1 Waddington (Wd)) whereas the tension corresponding to its daughters MS and E are 66.5 and 79.7 Wd, respectively. Furthermore, the daughters of E, which have committed to the endoderm fate demarcate a region on the landscape with an epigenetic tension of 130.6 and 143.1 Wd (Figure 1G and Table 1). An example of a rudimentary canal is that of the C lineage, which develops into body-wall muscle and hypodermis (Figure 1D,G). I consider the canals to be rudimentary since they do not resemble Waddington’s clear depressions with sides that prevent the metaphorical ball from exiting sideways. In the case of the AB, C and D lineages, it appears as if the ball could exit sideways and enter into the EMS lineage through the daughters of MS and E (Figure 1G). This apparent deficiency in the landscape could be interpreted biologically and technically. Biologically speaking, blastomeres of the early *C. elegans* embryo could indeed convert from one to another in the absence of necessary regulators. For example, the AB conversion to EMS blastomeres occurs with the loss of *mex-1* and *mex-5* function (<u>m</u>uscle in <u>ex</u>cess) and the C and D conversions occur with the loss of *pie-1* function (<u>p</u>harynx and intestine in <u>e</u>xcess) (Mello et al., 1992; Schubert et al., 2000). Technically, since epigenetic tension values are added equally to a given matrix region, this leads to regional flat surfaces. An alternative is to incorporate a Gaussian distribution factor to the epigenetic tension values added to a given region. Such a step would incorporate local curves within each region leading to borders that isolate each region from its neighbor. However, here I prefer to limit the generated surface to the actual calculated values with no visual embellishments other than those inherent in perspective rotations (Becker et al., 1988).

**Table 1.** Epigenetic tension and information entropy for *C.elegans* early embryogenesis

| Cell | Epigenetic Tension (Wd) | Shannon's Entropy (nat) |
| --- | --- | --- |
| P0 | n/a | 8.435443 |
| P1 | 9.37 | 8.401806 |
| P2 | 13.86 | 8.375532 |
| P3 | 25.30 | 8.189275 |
| P4 | 84.13 | 8.177531 |
| AB | 11.31 | 8.438601 |
| ABa | 20.85 | 8.39817 |
| ABp | 18.65 | 8.375853 |
| ABar | 62.86 | 8.247753 |
| ABal | 59.26 | 8.244755 |
| ABpr | 52.46 | 8.21 |
| ABpl | 72.10 | 8.297715 |
| ABalx | 66.57 | 8.109071 |
| ABarx | 62.90 | 8.076559 |
| ABplx | 65.09 | 8.0763 |
| ABprx | 73.76 | 8.042356 |
| EMS | 53.33 | 8.406477 |
| E | 79.74 | 8.175262 |
| Ea | 130.60 | 7.387439 |
| Ep | 143.12 | 7.845338 |
| MS | 66.50 | 7.585457 |
| MSx1 | 115.23 | 7.905293 |
| MSx2 | 100.08 | 7.846617 |
| C | 82.26 | 8.307016 |
| Cx1 | 97.57 | 7.792537 |
| Cx2 | 82.94 | 7.800279 |
| D | 48.38 | 7.757407 |

### Epigenetic tension correlates inversely with Shannon’s entropy

A Waddington landscape begins with pluripotency and ends with differentiation. A feature of pluripotent transcriptomes is global genome representation (a large portion of the genome is represented in the transcriptome). On the other hand, differentiated cells consolidate their transcriptomes and represent the genes required for function (Efroni et al., 2008; Percharde et al., 2017). A consequence of global genome representation during pluripotency is that the transcriptome has a higher level of information (Shannon’s) entropy (Shannon, 1948). Shannon’s entropy correlates inversely with epigenetic tension (Pearson’s correlation =−0.70, *p*-value = 6.573e-05) and in principle could also be used to compute a Waddington landscape for the *C. elegans* early embryo (**Supplemental Figure 1**). For instance, comparing the gene expression profiles from cells in the P1->EMS->E->Ep lineage demonstrates a clear shift in data distribution away from uniformity and a reduction in Shannon’s entropy (Figure 2A-B, Table 1, Methods). Although methods exist for entropy-based quantification of potency from single-cell RNAseq data (Banerji et al., 2013; Grün et al., 2016; Guo et al., 2017; Teschendorff and Enver, 2017), adopting epigenetic tension as the sum of upregulations has three advantages. First, the shift away from uniformity (the reduction in entropy) that occurs during blastomere specification is a result of a subset of genes being upregulated. In that sense, epigenetic tension captures the process leading to entropy reduction consistent with the inverse correlation between the two metrics. Second, all analyzed cells (except ABpl) had lower epigenetic tension than their daughters, but this consistency was clearly lost between MS and its daughters when calculating Shannon’s entropy (Table 1). While this discrepancy may be biologically relevant in the context of differentiation (see Discussion), epigenetic tension provided a more consistent metric. Lastly, I find that summing gene upregulations is more intuitive and consistent with Waddington’s rope-tension model and allows easy interrogation of genes contributing most to the final value (the most upregulated genes).

**Figure 2.**
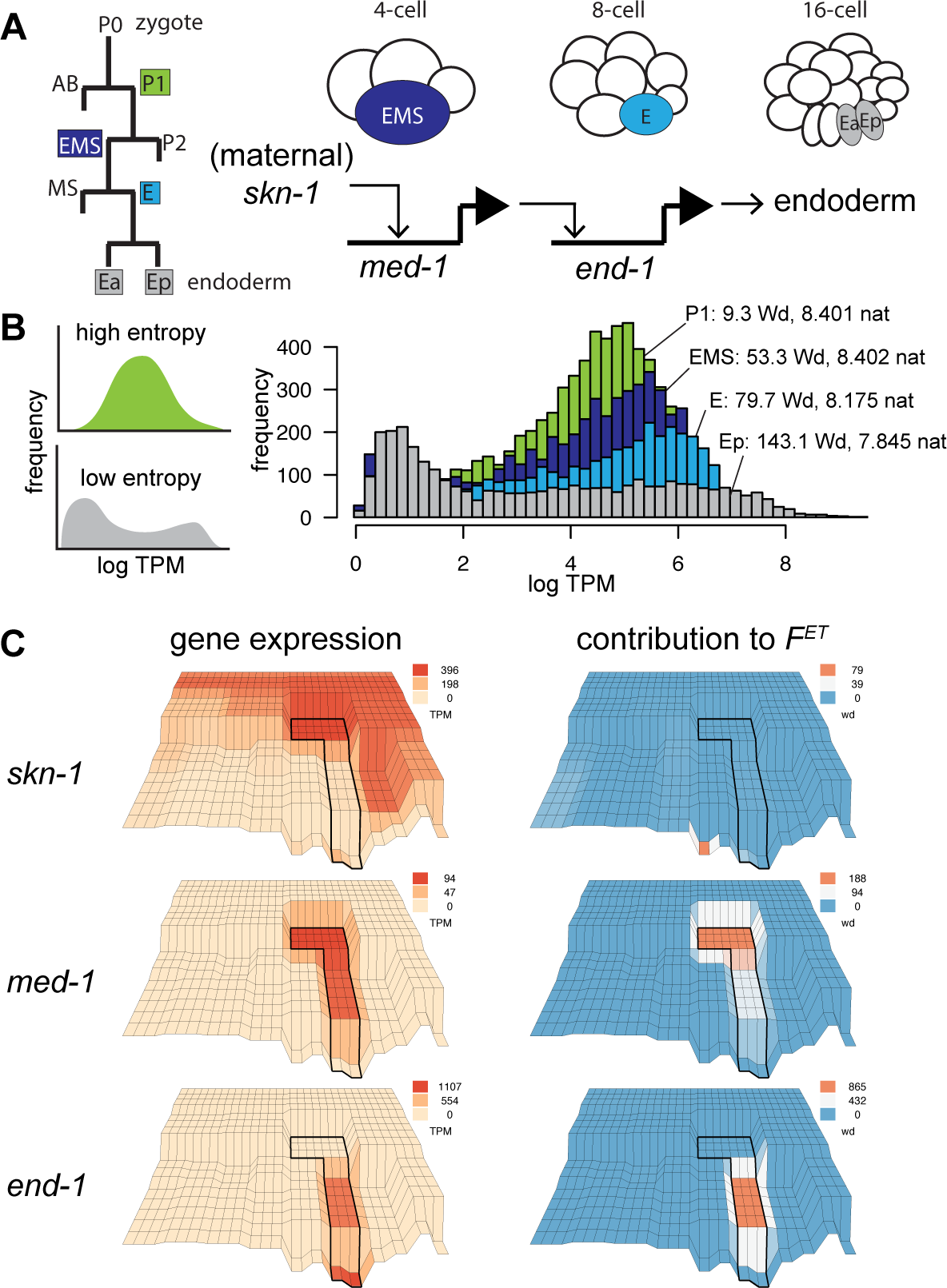
Reduced entropy and increased epigenetic tension during *C. elegans* endoderm specification. A) A brief overview of endoderm specification. The E lineage descends from P1 via EMS. Maternal *skn-1* induces *med-1* expression in EMS, which induces *end-1* expression in E to initiate endoderm differentiation. B) The pluripotent P1 transcriptome has higher information entropy (measured in nat) compared to differentiated Ep. Reduction in entropy correlates inversely with increase in epigenetic tension. C) Endoderm specification depicted on epigenetic landscape showing expression levels (left) and epigenetic tension (right). Note that *med-1* contributes to tension at EMS while *end-1* contributes to the tension at E. The lineage from EMS to E daughters is outlined. See Figure 1G for annotation.

### Endoderm specification as an example for interpreting the modeled landscape

The emergence of canals in the generated landscape is due to lineages having different degrees of gene upregulation leading to different epigenetic tensions. This can be demonstrating by tracing the origin of the canal leading to the worm intestine (Figure 2A). The endoderm lineage that builds the worm intestine originally emerges from a competent EMS blastomere where the transcription factor SKN-1 activates the endo-mesoderm specification factors *med-1/2*, which activate transcription factors *end-1/3* in E to initiate endoderm fate (Bowerman et al., 1992; Elewa et al., 2015; Lin et al., 1995; Maduro et al., 2015; Maduro et al., 2007) (Figure 2A). Inspecting the landscape, we see a sudden dip in the region corresponding to EMS (Figure 1G). The difference in epigenetic tension in this region is consistent with EMS being the only cell at the four-cell stage with considerable expression of new transcripts (Hashimshony et al., 2012). The dip at EMS coincides with maternal *skn-1* abundance (392.9 TPM) and zygotic *med-1* expression (94.2 TPM) (Figure 2C). The sudden spike in *med-1* expression contributes 188.4 wd to the epigenetic tension in this region (*med-2* contributes 237.2 TPM and 474.4 wd). This effect is propagated in the daughter cell E where *med-1* activates *end-1* expression (432.3 TPM), which contributes 864.6 wd to the epigenetic tension at E (*end-3* contributes 745.6 TPM and 572.3 wd). Taken together, a surface resembling the features of a Waddington Epigenetic Landscape can be created for early *C. elegans* embryogenesis by integrating the invariant cell lineage and sc-RNA data (Sulston et al., 1983; Tintori et al., 2016). Since the values leading to the topology are based on gene upregulation, the shape of the landscape is related to gene activity in agreement with the original intent of Waddington.

### Major contributors to the modeled landscape are at the nexus of stress response, behavior and transgene silencing

The depth at each region of the *C. elegans* epigenetic landscape is determined by the calculated epigenetic tension, which in turn is a function of gene upregulation. To explore the genes that contribute the most to shaping the modeled landscape, one can view the landscape from below and ask which genes contribute to the tension at each region (Figure 3A). For the sake of brevity, I will refer to genes that contribute to epigenetic tension as “tensors” and will refer to tensors in the 95% percentile (i.e. top 5% in contribution to epigenetic tension) as “top tensors” (**Supplemental File 1**). This threshold was chosen to minimize the effect of upregulation noise in subsequent analyses since no filters were used when calculating epigenetic tension from gene upregualtion. Phenotype enrichment analysis of the top tensors in each cell leading to the 16-cell embryo revealed that genes implicated in locomotion and nervous system function (behavior), stress response and transgene silencing shape the early embryo (Angeles-Albores et al., 2016). For example, the top tensors of ABarx were enriched for ectopic expression of transgene, dauer metabolism and muscle system phenotypes (Figure 3A). While a similar analysis for the tensors in E did not result in significant phenotype enrichments, the daughter cell Ea was enriched for phenotypes related to toxin hypersensitivity, fat content and response to acetylcholinesterase inhibition (Figure 3A). Combining together the top tensors from all 26 cells (4174 genes) confirms an enrichment in genes required for proper muscle cell and system morphology, regulating fat content and resistance to pore forming toxins (**Supplemental File 1**). A gene ontology analysis revealed an enrichment in genes involved in transcription regulation and metabolism, consistent with the phenotype enrichment analysis (**Supplemental File 1**).

**Figure 3.**
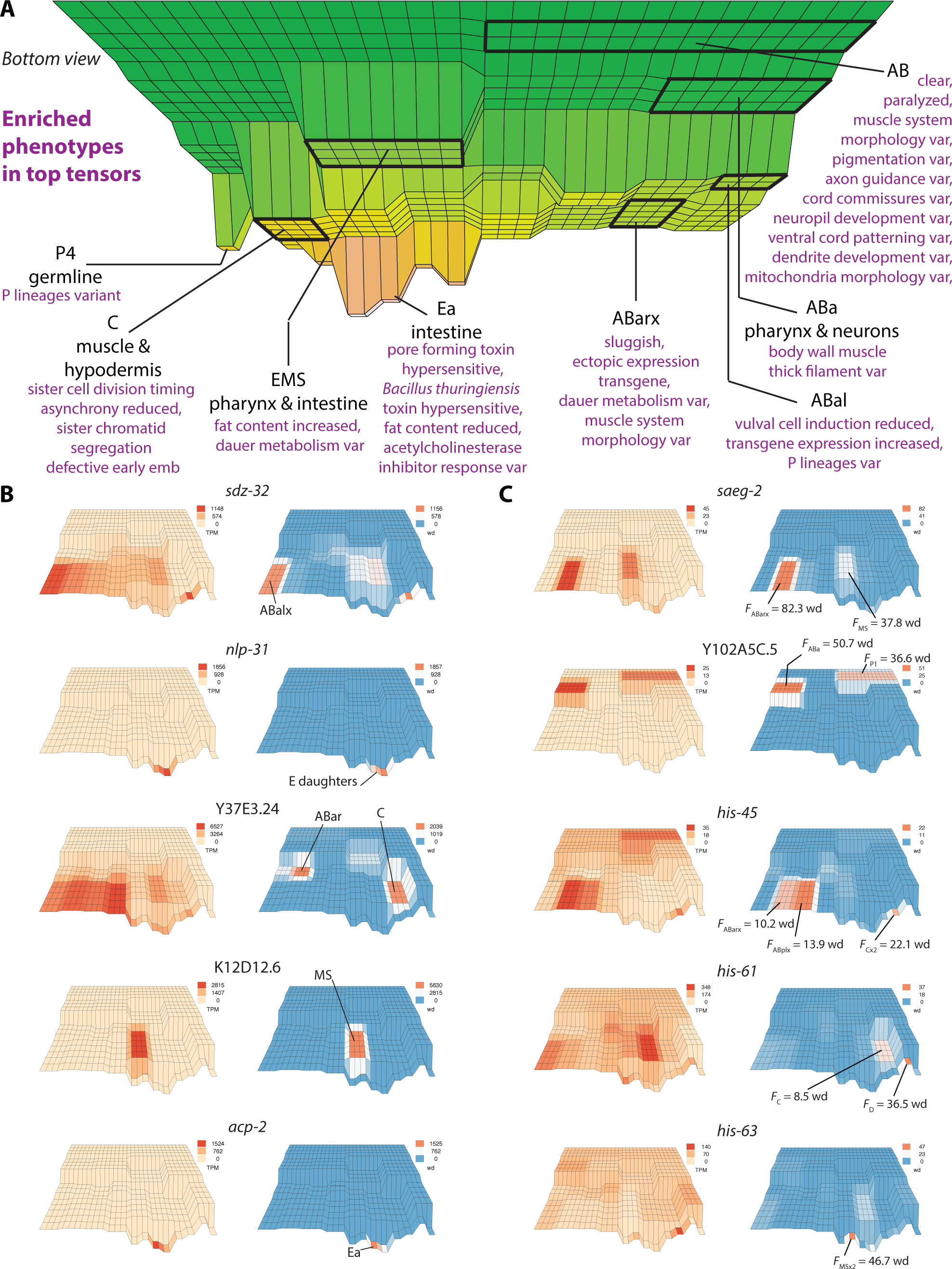
Stress and behavior shape the early epigenetic landscape. **A)** The view from under the landscape showing phenotypes enriched in top tensors for different cells (var = variant). **B)** Five examples of 39 tensors that contribute more than 1000 wd to the epigenetic landscape. **C)** Five genes that are regulated by neuronal small RNAs over multiple generations (Posner *et al*.) contribute to the early epigenetic landscape.

Thirty-nine of the top tensors contributed more than 1000 wd to the modeled landscape. Most of the 39 genes are either regulated by insulin receptor *daf-2* (e.g. *fbxb-81* and *sdz-32*), upregulated in dauer-constitutive mutants (e.g. C26F1.1) or affected by different forms of stress such as pathogen infection (e.g. *nlp-31*, *fipr-2* and *prx-3*) toxic chemicals (e.g. C29F7.2 and Y37E3.24) or high salt concentration (e.g. H05C05.4) (**Figure 3B, Supplemental Table 1 and Methods**). At the nexus between *daf-2* signaling and behavior is *nlp-12*, a *daf-2/daf-16*-dependent neuronal enriched gene (**Figure 3B**). In general, *daf-2/daf-16*-dependent neuronal genes are enriched for top tensors (*p*-value = 2.842e-11 Table 2) (Kaletsky et al., 2016). Interestingly, *acp-2* and Y65A5A.1 have been implicated in behavior and long-term memory, respectively (Cotella et al., 2013; Freytag et al., 2017). However, these two genes contribute more than 1000 wd to E daughter cells which build the worm intestine and do not contribute any neurons to the body. This observation will be elaborated upon below. In summary, genes with the highest contributions to epigenetic tension up to the 16-cell stage embryo straddle the nexus between behavior, transcription regulation, and various forms of stress response.

**Table 2.** Enrichment for top tensors in published datasets.

| Reference | Dataset | # Genes | Top tensors | adj. *p* value |
| --- | --- | --- | --- | --- |
| Kaletsky *et al.* | *daf-2/16* adult neuronal transcriptomes | 1058 | 221 | 2.842545e-11 |
| Moore *et al.* | Down in F1 of N2 PA14 fed | 3536 | 577 | 1.572315e-07 |
| Moore *et al.* | Up in F1 of N2 PA14 fed | 4805 | 810 | 6.960326e-14 |
| Moore *et al.* | Down in F2 of N2 PA14 fed | 228 | 29 | 1 |
| Moore *et al.* | Up in F2 of N2 PA14 fed | 1046 | 139 | 0.9233524 |
| Posner *et al.* | RDE-4 dependent NeuroSTGs | 46 | 18 | 0.0002159394 |
| Posner *et al.* | RDE-4 dependent GermSTGs | 124 | 30 | 0.00990078 |
| Posner *et al.* | RDE-4 dependent GermSTGs (whole worms) P0-F3 | 194 | 52 | 6.087601e-06 |
| Posner *et al.* | RDE-4 dependent GermSTGs Germline mRNA STG P0 | 40 | 9 | 1 |
| Vasale *et al.* | EGRO-1 targets | 60 | 25 | 1.056948e-06 |
| Gu *et al.* | WAGO-1 (IP) targets | 98 | 21 | 0.2126531 |
| Claycomb *et al.* | Down in *csr-1* mutant | 937 | 249 | 5.216495e-27 |
| Claycomb *et al.* | Down in *ego-1* mutant | 2125 | 388 | 1.206053e-10 |
| Claycomb *et al.* | CSR-1 (IP) | 4191 | 556 | 0.4866012 |
| Tu *et al.* | CSR-1 (IP) targets | 5070 | 753 | 0.000920504 |
| Tu *et al.* | Depleted in *csr-1* mutant | 2029 | 520 | 1.070983e-52 |
| Tu *et al.* | WAGO-1 (IP) targets | 1735 | 516 | 2.764214e-75 |
| Conine *et al.* | CSR-1 (IP) targets | 5574 | 799 | 0.01923358 |
| Conine *et al.* | ALG-3/4 (IP) targets | 1156 | 154 | 0.8789219 |
| Conine *et al.* | Down regulated in *alg-3/4* mutant | 214 | 23 | 1 |
| Conine *et al.* | Up regulated in *alg-3/4* mutant | 204 | 37 | 0.2706262 |
| Buckley *et al.* | HRDE-1 targets (strict) | 81 | 23 | 0.004201192 |
| Buckley *et al.* | HRDE-1 targets | 1587 | 436 | 2.41759e-52 |
| Bagijn *et al.* | PRG-1 targets (protein coding genes) | 13182 | 2107 | 3.486802e-18 |
| Bagijn *et al.* | PRG-1 targets (pseudogenes) | 759 | 78 | 0.04426326 |

### Transgenerational behavior genes contribute to the modeled landscape

The enrichment in genes that regulate transgene silencing, stress response and behavior among the top tensors of the early epigenetic landscape, suggested that early embryogenesis is shaped by the physiological adaptations of parents. Indeed, genes upregulated or downregulated in the progeny of animals that were fed pathogenic bacteria (PA14) were enriched in top tensors (*p*-value = 1.572e-07 and 6.960e-14, respectively **Table 2 and Methods**) (Moore et al., 2019). This was not the case for genes that continued to be regulated in the following generation (F2) (Table 2). However, it is likely that differential expression in F2 animals is a function of the specific experience encountered by grandparents and since the dataset used to generate the epigenetic landscape did not come from animals that were fed PA14, this discordance may be expected. A more appropriate analysis can be done by evaluating genes that are targeted by RDE-4-dependent transgenerational neuronal small RNAs that are generated while animals are maintained under normal laboratory conditions and fed the standard *E. coli* OP50 diet (Posner et al., 2019). Genes that are targeted by neuronal siRNAs (NeuroSTGs) and those targeted by neuronal siRNAs that are transmitted to the germline (GermSTGs) are significantly enriched for top tensors (18/46 and 30/124, respectively Table 2). Moreover, genes that continue to be targeted up to the fourth generation (F3) by RDE-4-dependent neuronal siRNAs are also significantly enriched for top tensors (52/194 Table 2). Posner *et al*., also identified forty genes that were differentially expressed in the germlines of animals with *rde-4(-)* neurons (i.e. not just targeted, but regulated by RDE-4 dependent GermSTGs). Nine of these 40 genes are top tensors (Table 2). Moreover, five genes continued to be regulated by GermSTGs until the F3 generation (i.e., upregulated in F3 of P0 animals with *rde-4(-)* neurons) including *saeg-2* a conserved gene that is required for foraging behavior (Hao et al., 2011). *saeg-2* is a top tensor of ABarx (82.36 wd) (Figure 3C). Another member of the Posner quintet is Y102A5C.5; a top tensor of two cells (50.7 wd in ABa and 36.6 wd in P1), and while the remaining three genes encoding for histones *his-45*, *his-61* and *his-63* are not among the 95^th^ percentile of tensors, unlike several others (i.e. *his-1/3/4/24/31/38*), they do contribute to the early epigenetic landscape (Figure 3C).

If transgenerational behavior shapes the early epigenetic landscape, then top tensors should also be targeted by small RNA pathways involving PRG-1, WAGO-1, HRDE-1, and/or CSR-1 (Bagijn et al., 2012; Buckley et al., 2012; Claycomb et al., 2009; Gu et al., 2009; Tu et al., 2015). Indeed, small RNAs binding to these argonautes, or dependent on their activity, target genes that are significantly enriched for top tensors (Table 2). However, immunoprecipitated targets of ALG-3/4, paralogous argonautes expressed during male gametogenesis and required for male fertility, are not significantly enriched for top tensors, nor are genes that are regulated by *alg-3/4* function (Table 2). Finally, a distinct class of small RNAs (26Gs) interact with the argonaute ERGO-1 and target genes expressed in the early *C. elegans* embryo (Vasale et al., 2010). While the fact that *ergo-1* mutants exhibit no developmental phenotypes has left the biological significance of ERGO-1-dependent 26Gs unresolved, ERGO-1 targets are significantly enriched in top tensors (25/60) suggesting a role for this argonaute in shaping the early epigenetic landscape (Table 2). Taken together, the top tensors shaping the modeled landscape are enriched for genes targeted by small RNA pathways including genes that propagate transgenerational behaviors.

### Several genes shape one lineage and later function in another

Curiously, performing an analysis for tissue expression enrichment revealed that many top tensors shape the beginning of one lineage even though their reported function or enrichment is in another lineage (Figure 4A-E and **Supplemental File 1**) (Angeles-Albores et al., 2016; Murray et al., 2012). Two examples referred to above are *acp-2* and Y65A5A.1, genes implicated in long-term memory but that contribute more than 1000 wd to the epigenetic tension of E daughter cells. Conversely, *elt-7* is a transcription factor involved in endoderm differentiation (Dineen et al., 2018; Maduro et al., 2015; Sommermann et al., 2010) but is a top tensor for the ABalx cell, which contributes neurons, hypodermis and the anterior pharynx. Another example is *ref-1*, which encodes a basic helix–loop–helix (bHLH) protein that antagonizes neurogenesis and is upregulated in response to Notch signaling in both the AB and E lineages in the embryo (Neves and Priess, 2005). However, *ref-1* is a top tensor for the C blastomere, even though it has not been reported to influence the development of this lineage. Two additional examples are *capa-1* and *tsct-1*; genes enriched in neurons under *daf-2/daf-16* regulation (Kaletsky et al., 2016). *capa-1* encodes an insect neuropeptide related gene that exhibits G protein-coupled receptor binding activity. Insect CAPA is related to vertebrate Neuromedin U (NMU) a structurally highly conserved neuropeptide of which highest levels are found in the pituitary and gastrointestinal tract (Lindemans et al., 2009). *tsct-1* is an ortholog of human TSC22D3 (TSC22 domain family member 3) and is predicted to have DNA-binding transcription factor activity (Kim et al., 2018). Though both genes are neuronally enriched, they are top tensors of the E blastomere. A final example is *ilys-3*, which is an gene with enriched expression in the intestine but is a top tensor of the ABplx cell, which gives rise to neurons and hypodermis (McGhee et al., 2007). In summary, several genes shape the modeled landscape at lineages different than the ones that they later on differentiate, pattern or in which they are highly expressed.

**Figure 4.**
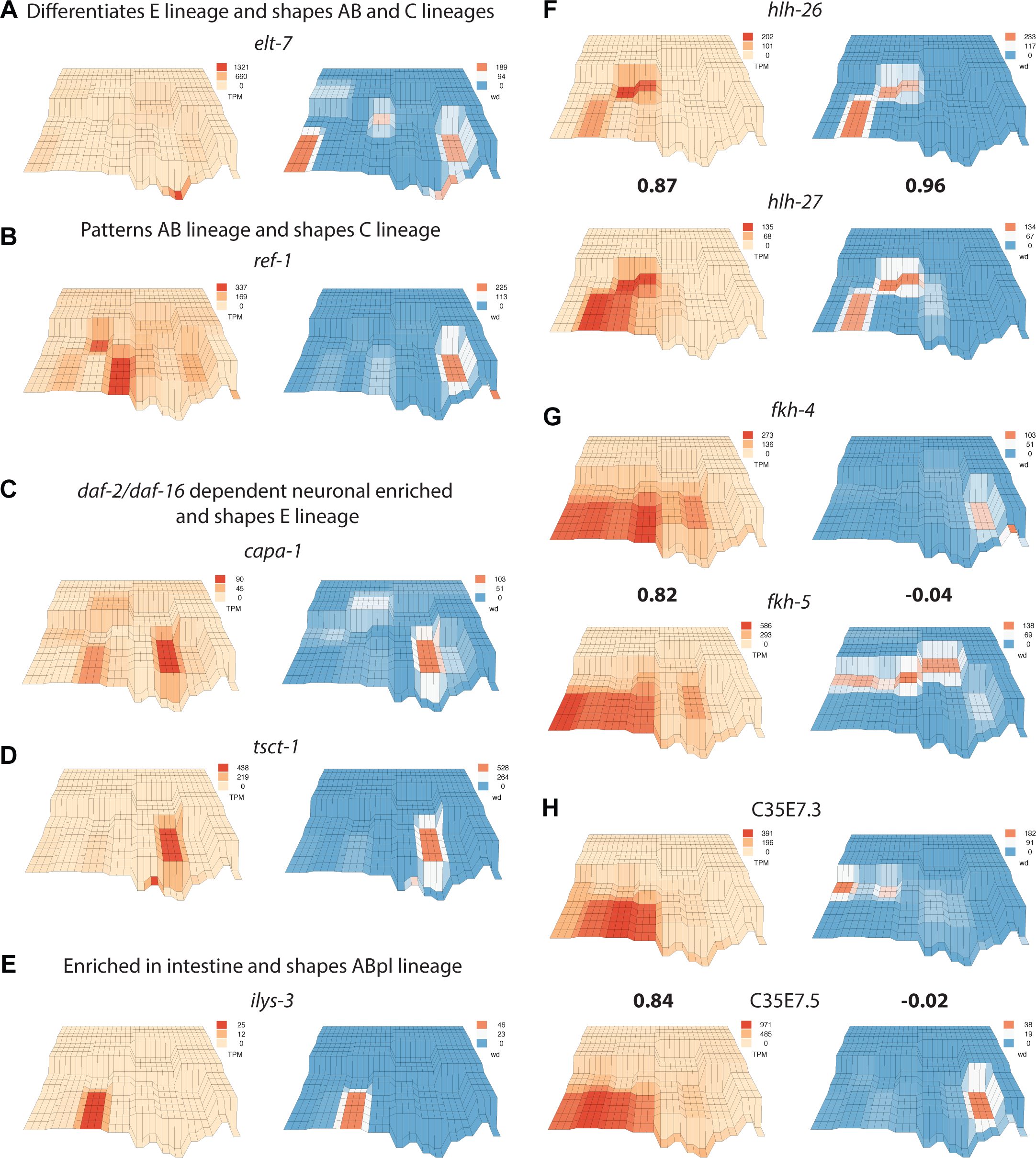
The disparity between expression profiles and contribution to epigenetic tension. **A)** *elt-7* is expressed in E daughter cells (>1000 TPM) but shapes the landscape of AB and C lineages. **B)** *ref-1* patterns AB granddaughters and is expressed in ABprx, but contributes to the epigenetic tension at C. **C-D)**. *capa-1* and *tsct-1* are enriched in neurons but are expressed in and contribute to the epigenetic tension at E. **E)** Expression of *ilys-3* is enriched in adult intestine but embryonically is enriched in and contributes to epigenetic tension of ABplx. **F)** *hlh-26* and *hlh-27* correlate in expression profile and contribution to epigenetic tension. **G-H)** *fkh-4/5* and C35E7.3/5 each correlate in expression profile but not in contribution to epigenetic tension. Pearson’s correlation in bold.

### Paralogs with similar expression profiles usually have different tension profiles

A third of *C. elegans* genes are predicted to have one or more paralogs and are therefore an ample source for genetic redundancy (Woollard, 2005). Paralogous genes expressed in the early *C. elegans* embryo are more likely to have similar expression profiles (i.e. synexpressed) compared to non-paralogous pairs (Tintori et al., 2016). However, the correlation in gene expression between paralogs is not coupled with a correlation in contribution to epigenetic tension. For example, while paralogs *hlh-25/26/27/28/29* have similar expression profiles and contributions to epigenetic tension this is not the case for most other synexpressed paralogs (**Figure 4F-H, Supplemental File 1**). Paralogous Forkhead transcription factors *fkh-3*, *fkh-4* and *fkh-5* have highly correlated expression profiles in anterior AB great-granddaughters and the E blastomere. However, *fkh-4* contributes to the epigenetic tension at C and *fkh-5* contributes mostly to EMS and the granddaughters of AB (**Figure 4G and Supplemental File 1**). Likewise, C35E7.3, C35E7.5 are highly correlated in their expression in AB great-granddaughters but have completely different contributions to epigenetic tension (Figure 4H). In summary, contribution to epigenetic tension, which is a function of upregulation, is not correlated to gene expression profile, meaning that synexpressed paralogs that are redundant in sequence and expression profile usually contribute to epigenetic tension differently.

## Discussion

Within an embryo intersects ancient laws of genesis and the ephemeral spirit of times. The two threads intertwine, the later upon former, and bring forth a fixed yet malleable being. Which parts of development fall within the domain of genetics, and which are influenced by recent history? This work offers the following insight: at the nativity of embryonic lineages, *C. elegans* blastomeres are epigenetically shaped by parental adaptations. So, while blastomere specification and subsequent differentiation may rely on conserved and robust genetic programs, they are hued by yesterday. This insight is a result of generating and exploring a Waddington epigenetic landscape. The generated landscape is a function of gene upregulation as determined from single-cell RNAseq data (Tintori et al., 2016). Therefore, the landscape does not capture other epigenetic dimensions related to post-transcriptional and post-translational regulation. Nevertheless, the landscape is based on a metric that, when plotted on a two-dimensional surface resembling the embryonic lineage, leads to a downward tilt with emerging canals. This metric is epigenetic tension, the sum of gene upregulations, and is highly influenced by top tensors, genes that are rapidly upregulated relative to their status in mother and sister cells.

What is an epigenetic tensor? The transcriptome of a pluripotent cell is more random than that of a differentiated cell and differentiation entails a reduction in the portion of the transcribed genome, which reduces its information entropy (Banerji et al., 2013; Efroni et al., 2008; Grün et al., 2016; Percharde et al., 2017). This reduction in entropy (and increase in order) occurs when fewer genes occupy more of transcriptome space, which is a consequence gene upregulation. When a gene rises from 20 TPM to 2000 TPM, it occupies more parts of the transcriptome and contributes to a reduction in entropy and an increase in differentiated order. In that sense, an epigenetic tensor is a gene that makes a significant contribution to transcriptome order in its trajectory towards differentiation. However, a simple thought experiment reveals that the opposite is also true. A differentiated cell undergoing reprogramming begins to engage in hypertranscription leading to a more global genome representation (Percharde et al., 2017). This means that more genes become upregulated, leading to an increase in epigenetic tension as well, but in the direction towards increased entropy, not order. The common factor in both scenarios is that an epigenetic tensor is a gene that is upregulated during transitions in cell identity.

Using epigenetic tension to generate a Waddington landscape imposes an interpretation of the landscape whereby differentiation and reprogramming are both downhill trajectories and where terminal maintenance or steady-state pluripotency (as with stem cells and oocytes) are elevated plateaus with few if any upregulated genes. Previous interpretations of Waddington’s landscape to convey cellular reprogramming depict a ball moving backwards uphill (induced pluripotency) or sideways over barriers to neighboring canals (transdifferentiation) (Hochedlinger and Plath, 2009; Yamanaka, 2009) (**Figure 5A**). However, according to my interpretation of the model, such depictions do not acknowledge developmental time. For a ball to move uphill a Waddington landscape is akin to saying that the developing system went back in time since “*the path followed by the ball as it rolls down toward the spectator, corresponds to the developmental history*” (Waddington, 1957). Likewise, to consider transdifferentiation a sideways move across the landscape does not incorporate developmental time in the transition from one cell identity to another. While canalization emphasizes that a metaphorical ball rolling down a hill is resistant to perturbations pushing it sideways to neighboring canals, it does not preclude the momentum of a ball allowing an ascent towards a steady-state plateau (Figure 5B). There, the ball (i.e. developing system) is at rest but is ready for activation. In the case of a *C. elegans* oocyte, the system is at a plateau of pluripotency ready for a fertilizing sperm, which would initiate the downhill roll of embryonic development. Here, I acknowledge, that by incorporating ramps that lead to *elevated* steady-states I may be parting from Waddington’s original conception of the landscape where “*the slope of the valley flattens out as it reaches the adult steady state*” (Waddington, 1957).

**Figure 5.**
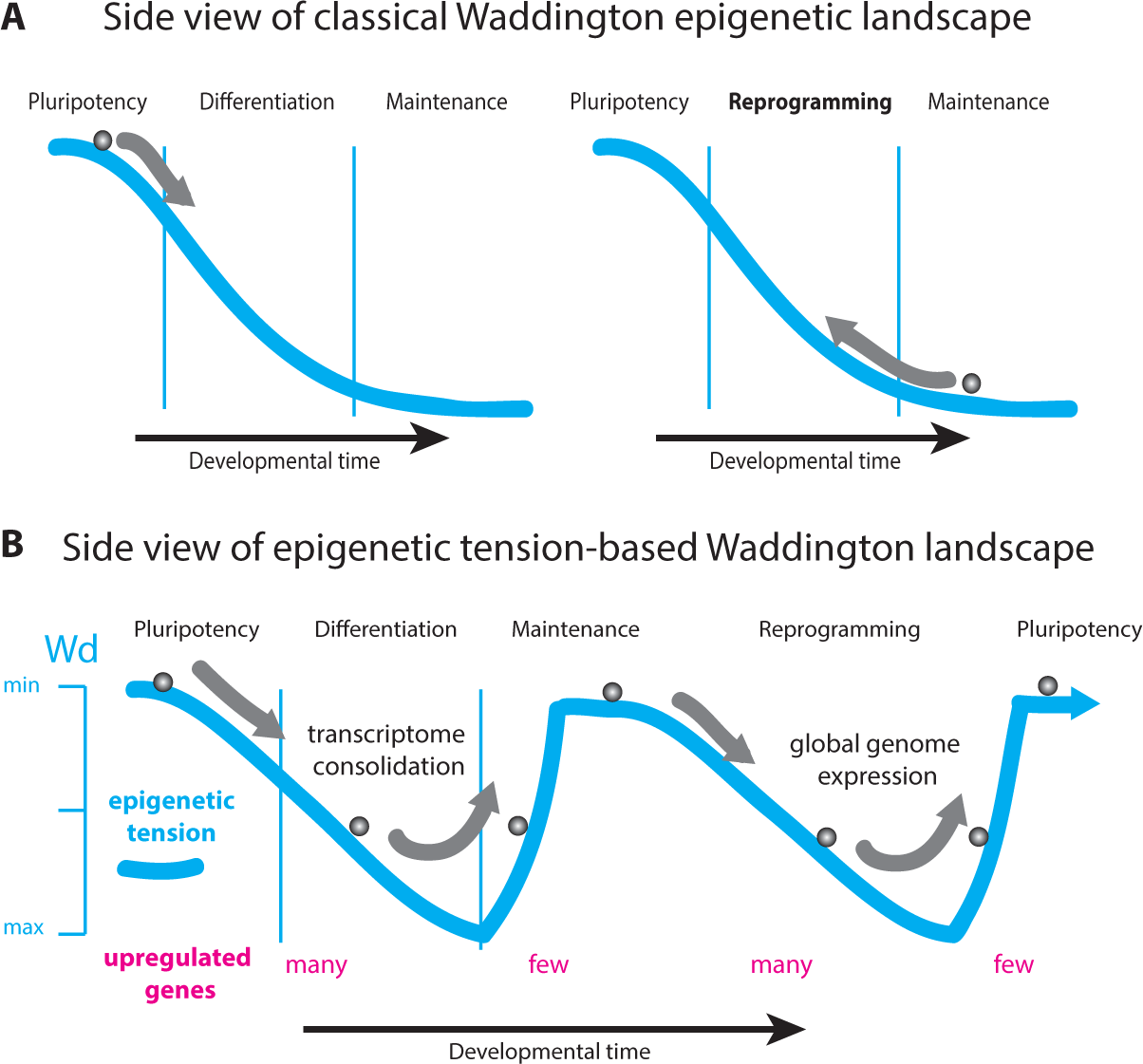
Adding momentum to Waddington’s Epigenetic Landscape. **A)** A side view of Waddington’s landscape and a common depiction of cellular reprogramming akin to going back in developmental time. B) An alternative interpretation of Waddington’s landscape imposed by the definition of epigenetic tension. Note that both pluripotency and terminal maintenance of a differentiated state are elevated plateaus. This would require allowing the ball to ascend to the steady state with the momentum of its descent.

Examining top tensors of the generated landscape suggests that transgenerational adaptations shape *C. elegans* embryogenesis. How could adaptations shape robust embryo development? Many of the top genes shaping the landscape encode proteins with F-box domains, known for their role in facilitating protein/protein interactions (Kipreos and Pagano, 2000) (**Supplemental File 1**). The *C. elegans* genome encodes hundreds of F-box genes, of types A, B and C. This expansion in the nematode repertoire of F-box genes is an outlier among animals (Kipreos and Pagano, 2000). It is conceivable that F-box genes in their plentitude offer an interphase between transgenerational cues and the network of protein interactions within a nascent blastomere. For example, a subset of F-box genes can be upregulated in response to small RNAs conveying a particular parental experience. This differential abundance in F-box proteins could then affect the protein interactome during blastomere specification and thereby tune the blastomeres and their lineages to resonate with the anticipated world. A similar effect can be envisaged for the multitude of histone genes in *C. elegans*. Three histones genes (*his-45*, *his-61* and *his-63*) are regulated for multiple generations by RDE-4-dependent small RNAs generated in neurons (Posner et al., 2019) and several other histone genes are top tensors (**Supplemental File 1**). It is conceivable that the variety of histones present during blastomere specification could be tuned by parental cues and define an adaptable context for robust genetic programs. F-box proteins and histones are therefore examples of how redundancy can serve as an interphase between adaptation and robustness during embryo development.

*saeg-2* is another gene regulated for multiple generations by RDE-4-dependent small RNAs generated in neurons (Posner et al., 2019). Importantly, this *saeg-2* transgenerational regulation was observed in the germline, not in neurons. Since the HRDE-1, the argonaute responsible for heritable silencing, is also restricted to the germline, this has led Posner *et al*. to suggest that neuronal RDE-4 influences behavior by regulating germline genes. Given the contribution of *saeg-2* to the generated landscape and since targets of HRDE-1 are enriched for top tensors, a possible mechanism for this germline-to-neuronal regulation is that neuronal RDE-4-dependent small RNAs transmitted through the germline shape the blastomeres that give rise to the nervous system and impact their overall responsiveness to categories of stimuli. Another possible mechanism is hinted at by the paralogous Forkhead transcription factors *fkh-3/4/5* (Figure 4G). *fkh-3/4/5* bind to promoters upstream of *C. elegans* piRNAs and promote their transcription by RNA Pol II (Cecere et al., 2012). Since piRNAs are detected in neurons from Aplysia to mammals (Lee et al., 2011; Rajasethupathy et al., 2012). It is possible that *fkh-3/4/5* shape the embryonic epigenetic landscape by promoting the expression of somatic piRNAs that relay particular silencing cues to neurons.

While *fkh-3/4/5* share a similar expression profile in the early embryo, their upregulation magnitudes differ significantly, leading to different contributions to epigenetic tension (Figure 4G). A related disparity is that observed with *elt-7* and *ref-1*, whereby genes shape the epigenetic landscape for one lineage but then differentiate, pattern or function in another (Figure 4A-B). This “entanglement” of lineages may be a way for lineages to inform one another of their configurations so that they develop into compatible organs. For example, an embryonic nervous system that is being informed of past starvation may develop more sensitive to food stimuli and would benefit from the cooperation of a digestive system that is more efficient in metabolism. Likewise, transmitting the memory of an encounter with harmful food could initialize neuronal and endodermal lineages so that they respond coherently to an otherwise attractive stimulus. The physiological communication between developed organs may have an embryonic form that exists during blastomere specification, whereby regulators of one lineage shape the beginning of another. Recall *capa-1*, a *daf-2/daf-16*-dependent neuronal enriched gene that contributes to the epigenetic tension of endoderm blastomere E (Figure 4C). *capa-1* is related to vertebrate neuropeptide Neuromedin U, which is expressed at high levels in the pituitary and gastrointestinal tract (Lindemans et al., 2009). Perhaps *capa-1* connects the functions of the worm intestine and neurons through its early shaping of the former and later enrichment in the second.

In closing, generating and exploring a Waddington epigenetic landscape for *C. elegans* supports an emerging theme that embryogenesis begins with a deposit of parental RNA that not only ensures robust development, but a development that is adapted to the world to come. •

## Acknowledgements

I thank Benjamin Roche and Ammar Tareen for valuable comments on the manuscript. This work was supported by NY State Department of Labor Unemployment Benefits and the patronage of Mostafa Nashaat.

## Methods

### Epigenetic Tension

Epigenetic tension was calculated from Tintori *et al*. Supplemental Table 2. First, RPKM values were scaled to TPM and then gene expression was averaged across replicates of each of the 27 embryonic cells. Note that in this dataset, daughters of ABal, ABar, ABpl, ABpr, MS and C are ambiguous and are represented by one cell for each pair (e.g. ABalx instead of ABala and ABalp).

Trimmed mean of M-values (TMM) normalization (Robinson and Oshlack, 2010) was not adopted for the following two reasons. First, the Tintori *et al*. dataset is based on single-cell RNAseq and TMM normalization is designed with bulk RNAseq in mind. Second, even when using the mean TPM values of replicates for each cell, which reduces zero-inflation and makes the data closer to bulk RNAseq, TMM normalization is not suitable since it was designed with a preliminary assumption that is not satisfied with the Tintori *et al*. dataset. Namely, TMM normalization adjusts for the composition of the RNA population that is being sampled and is recommended when the majority of the genes are believed not differentially expressed between any pair of the samples. Epigenetic tension relies heavily on the difference in RNA composition, which transitions from highly diverse during pluripotency to more homogenous as the transcriptome approaches differentiation (Figure 2B). Therefore, TMM normalization would dampen the very effect epigenetic tension attempts to highlight. One may then argue that TMM normalization could be used for replicates of the same cell type to adjust for variation in RNA composition. But this would mean applying TMM normalization to zero-inflated single-cell RNAseq data, which is problematic.

To calculate epigenetic tension, the expression of each gene in a cell is divided by its expression in its mother cell and added to the ratio of its expression relative to its sister cell. Only ratios greater than 1 (i.e. upregulation) are included in the calculation. Ratios equal to or less than 1 are converted to 0. An example for the calculation conducted in R for blastomere EMS is as follows:

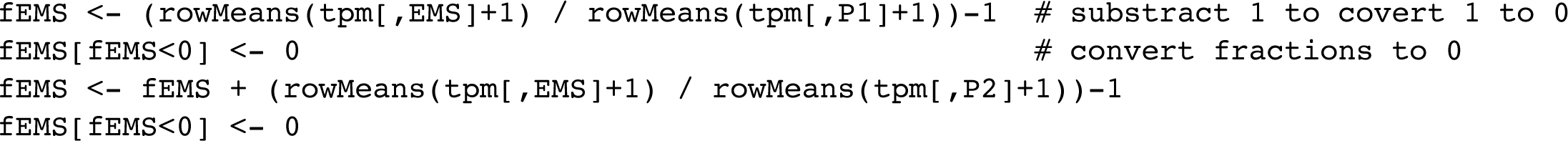

The result of this calculation is 26 epigenetic tensions corresponding to the 27 embryonic cells except for P0 (zygote). Since P0 has no mother or sister the calculation described above cannot be performed. For the the purpose of generating an epigenetic landscape, the value give to P0 was the mean of the epigenetic tension values for its daughters AB and P1.

### Shannon’s Entropy

Shannon’s entropy was calculated as

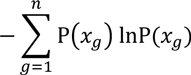

where *x* is gene expression in TPM for each gene (*g*). The following calculation in R was applied to the averaged gene expression values for each of the 27 cells (*i*).

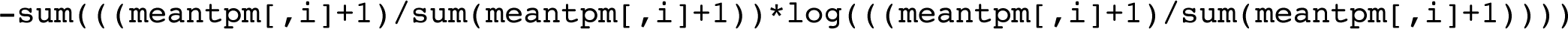

### Phenotype, Tissue expression and Gene ontology enrichment

Phenotype, Tissue expression and Gene ontology enrichment analysis was performed using the gene set enrichment tool provided by Wormbase. https://wormbase.org/tools/enrichment/tea/tea.cgi

### Top tensor expression and enrichment

Expression data for the 39 top tensors contributing more than 1000 wd (**Supplemental Table 1**) was curated from Wormbase “Expression Cluster” data and reference therein. Enrichment for the 4174 top tensors in published datasets (Table 2) was tested using a hypergeometic distribution after a Bonferroni correction for an adjusted *p*-value.

---

**Supplemental Figure 1.**
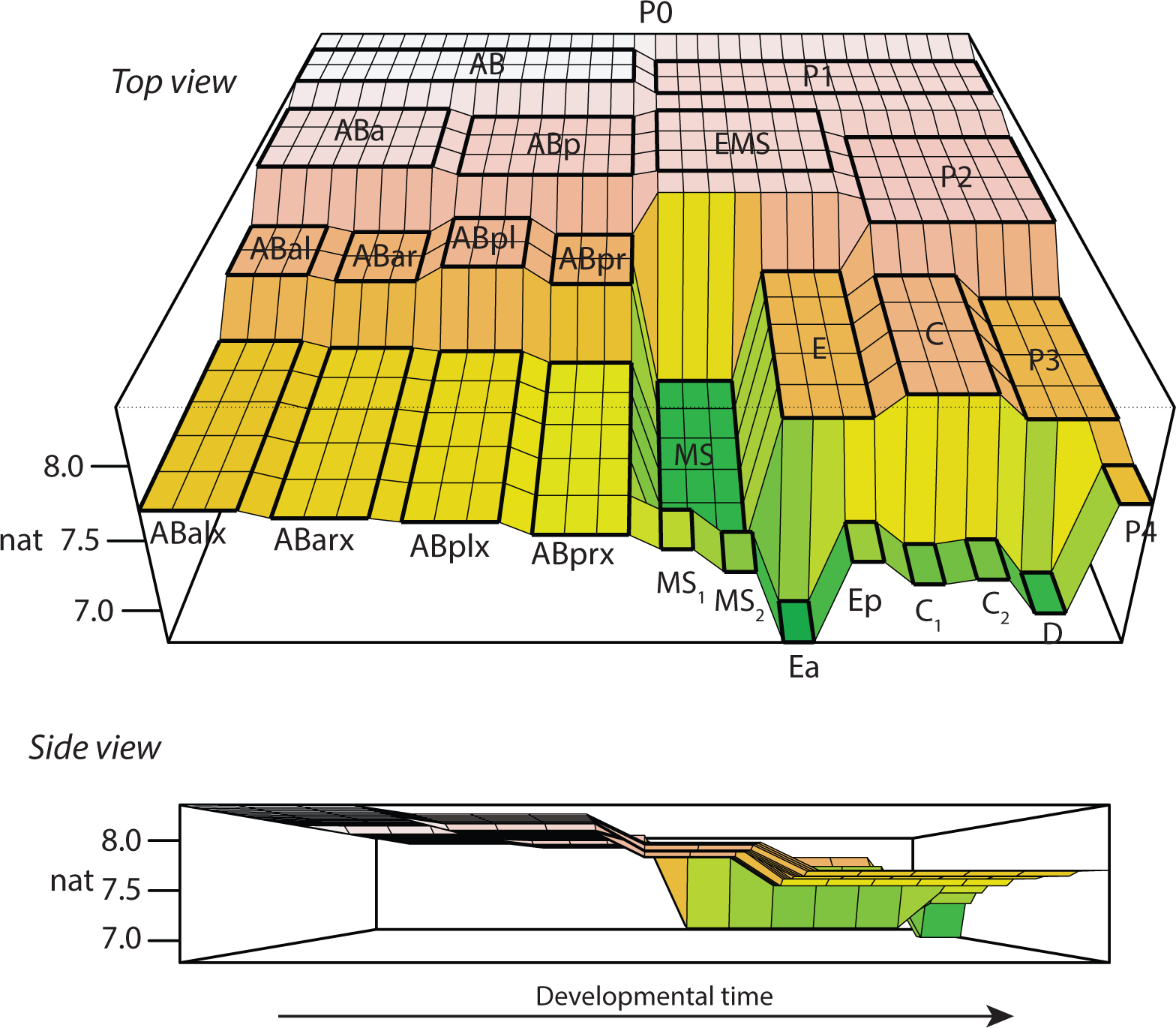
A Waddington epigenetic landscape based on Shannon’s entropy. A regionalized matrix representing the C. elegans early embryo lineage is populated with the information entropy value for each cell based on the Tintori *et al*. dataset. Note the rise in MS daughter cells compared to their mother.

**Supplemental Table 1:** Top tensors contributing more than 1000 wd are at the nexus of Aging/Stress and Behavior

| Cosmid no | Gene | Description | Cell | F<sup>ET</sup>(wd) | Aging/Stress | Behavior |
| --- | --- | --- | --- | --- | --- | --- |
| AH10.2 | - | - | EMS | 1064.8 | 5, 11-13 | 7, 14 |
| B0213.6 | *nlp-31* | Neuropeptide-like protein | Ea, Ep | 1182.8, 1856.8 | 11, 17, 19, 25, 32-34, 40, 42-45 | 10, 28 |
| C15H9.8 | *prx-3* | Peroxisome assembly factor | Ea, Ep | 1148, 1093.3 | 11, 12, 19, 21, 22, 32, 35, 48, 49 | 7, 27, 41 |
| C26F1.1 | - | - | Ea | 1631 | 2, 4, 6, 11, 13, 15-19 | 7, 10 |
| C29F7.2 | - | Protein kinase-like domain | Ea | 1772.4 | 1, 11, 12, 15, 19-26 | 7, 27, 28 |
| C52E2.7 | *fbxb-96* | F-box B protein | Ea | 3005.9 | 5, 11-13, 23, 32, 33 | 7 |
| F09C3.3 | *fbxb-68* | F-box B protein | D | 1162.7 | 5, 13, 35 | 7, |
| F11C1.2 | *mff-1* | Mitochondrial Fission Factor | Cx1 | 1105.1 | 34, 4, 12 | 7, 28, 10, 41 |
| F14E5.4 | *acp-2* | Acid phosphatase family | Ea | 1525 | 1-6 | 7-10 |
| F23H12.9 | *fipr-2* | Neuropeptide-like protein | Cx1 | 2227.5 | 2, 4, 6, 11-13, 16, 17, 19, 25, 39, 40 | 28 |
| F31E9.10 | - | Pseudogene | Cx1 | 1018.9 | - | - |
| F37D6.4 | - | F-box associated domain | ABarx | 1084.5 | 5, 11-13, 23, 32, 33 | 7 |
| F40G9.20 | - | ncRNA | ABalx, ABplx, MSx2 | 1169.8, 1458.6, 1046.2 | 25 | - |
| F52D2.8 | *fbxa-46* | Pseudogene | Ep | 1004 | 4, 12, 13, 33, 34 | 7 |
| F58G4.4 | *sdz-23* | SKN-1 dependent zygotic | E | 1364.8 | 11, 12, 13 | 7 |
| F59D12.11 | - | ncRNA | MS | 1173.6 | - | - |
| H05C05.4 | - | SUP-like | Cx1 | 3587.5 | 5, 19, 25 | 10, 41 |
| K10C9.9 |  | Pseudogene | Ea | 1409 | 11, 12, 25, 33 | 10 |
| K12D12.6 | - | ncRNA | MS | 5629.5 | - |  |
| M01D7.5 | *nlp-12* | Neuropeptide-like protein | Cx2 | 1139.6 | 4, 11 12, 17, 19, 24, 25, 26, 29, 37, 40, 46, 47 | 28, 41 |
| R07C3.9 | *fbxc-31* | F-box C protein | D | 1832.1 | 4, 11-13, 35 | 7 |
| T04D1.2 | - | F-box associated domain type 2 | D | 1086.7 | 4, 5, 11-13, 19, 32, 39 | 14, 28 |
| T05E12.10 | - | ncRNA | C, EMS, P4 | 2790.2, 1411.9, 1504.1 | 13 | 50 |
| T07D1.8 | - | ncRNA | Ea | 1156.3 | - | - |
| T09B4.4 | - | Calmodulin like 4 | Ea | 2981 | 2, 11, 25 | 7, 10, 28, 38 |
| T10E9.8 | - | - | D | 1592.6 | 5, 11, 12 | 7, 28, 10 |
| T17A3.3 | *fbxb-81* | F-box B protein | Cx1 |  | 4, 5, 11-13, 23, 35 | 7 |
| Y23H5B.1 | - | Domain of unknown function | Ep | 1200.4 | 5, 11 | 7, 28 |
| Y37E3.24 | - | ncRNA | ABar, C | 2038.6, 1916.6 | 19 | - |
| Y37E3.25 | - | ncRNA | ABar, C | 1009.6, 1615.5 | - | - |
| Y40B1B.3 | *fbxb-66* | F-box B protein | D | 1135.6 | 4, 5, 11-13, 19, 23, 35-37 | 7, 38 |
| Y48G8AL.15 | - | Nucleoside diphosphate kinase | D | 1081.5 | 4, 5, 12, 19, 22, 25, 29, 51 | 7, 28 |
| Y49F6B.5 | *sdz-32* | SKN-1 dependent zygotic | Cx2 | 1156.3 | 11-13, 34 | 7 |
| Y65A5A.1 | - | - | Ea | 1804.9 | 11-13, 17, 19, 24, 29, 32, 35, 52, 53 | 7, 28, 38 |
| Y75B12A.2 | - | - | EMS | 1030.8 | 2, 4, 5, 11-13 47 | 7 |
| Y92H12BR.4 | - | Pseudogene | Ea | 4082.3 | 4, 11, 12, 29 | 54 |
| Y97E10AM.1 | *ceh-76* | C. elegans homeobox | MSx1, MSx2 | 1216.7, 1795.8 | 4, 12, 13, 19, 33 | 7 |
| ZK1193.5 | *dve-1* | SATB homeobox 2 | Ea, Ep | 1047.6, 1175 | 16, 29, 30 | 28, 31 |
| ZK185.3 | - | Predicted glycine N-acyltransferase | Ea | 1368.9 | 11, 12, 19, 21, 22, 29, 32, 47, 53 | 7, 28, 41 |

